# A Machine Learning Enhanced EMS Mutagenesis Probability Map for Efficient Identification of Causal Mutations in *Caenorhabditis elegans*

**DOI:** 10.1101/2024.02.15.580605

**Authors:** Zhengyang Guo, Shimin Wang, Yang Wang, Zi Wang, Guangshuo Ou

## Abstract

Chemical mutagenesis-driven forward genetic screens are pivotal in unveiling gene functions, yet identifying causal mutations behind phenotypes remains laborious, hindering their high-throughput application. Here, we reveal a non-uniform mutation rate caused by Ethyl Methane Sulfonate (EMS) mutagenesis in the *C. elegans* genome, indicating that mutation frequency is influenced by proximate sequence context and chromatin status. Leveraging these factors, we developed a Machine Learning enhanced pipeline to create a comprehensive EMS mutagenesis probability map for the *C. elegans* genome. This map operates on the principle that causative mutations are enriched in genetic screens targeting specific phenotypes among random mutations. Applying this map to Whole Genome Sequencing (WGS) data of genetic suppressors that rescue a *C. elegans* ciliary kinesin mutant, we successfully pinpointed causal mutations without generating recombinant inbred lines. This methodology can be adapted in other species, offering a scalable approach for identifying causal genes and revitalizing the effectiveness of forward genetic screens.

**Significance statement:** Exploring gene functions through chemical mutagenesis-driven genetic screens is pivotal, yet the cumbersome task of identifying causative mutations remains a bottleneck, limiting their high-throughput potential. In this investigation, we uncovered a non-uniform mutation pattern induced by Ethyl Methane Sulfonate (EMS) mutagenesis in the *C. elegans* genome, highlighting the influence of proximate sequence context and chromatin status on mutation frequency. Leveraging these insights, we engineered a machine learning enhanced pipeline to construct a comprehensive EMS mutagenesis probability map for the *C. elegans* genome. This map operates on the principle that causative mutations are selectively enriched in genetic screens targeting specific phenotypes amid the backdrop of random mutations.

Applying this mapping tool to Whole Genome Sequencing (WGS) data derived from genetic suppressors rescuing a *C. elegans* ciliary kinesin mutant, we achieved precise identification of causal mutations without resorting to the conventional generation of recombinant inbred lines. Our work not only advances understanding of mutation dynamics but also revitalizes the efficacy of forward genetic screens, contributing to the refinement of genetic exploration methodologies with implications for various organisms.

## Introduction

Forward genetic screens have been instrumental in demystifying the molecular mechanisms underpinning myriad biological processes across a diverse array of organisms. The procedure commences with the creation of a genetically varied population, often achieved via the application of chemical mutagens such as Ethyl Methane Sulfonate (EMS) (1–3) or through irradiation (4, 5), thereby seeding the genome with a variety of mutations. This random mutagenesis generates a pool of organisms each harboring a distinct set of genetic aberrations. Subsequent stages involve diligent phenotypic screening, identifying responsible genes, and decoding biological significance. This strategy has been pivotal in unveiling gene functions across multiple biological spectra, spanning developmental processes (6, 7), signal transduction (8), and disease mechanisms (9). Genes unearthed through these screens often illuminate deeper molecular mechanisms and are frequently employed to glean insights into analogous processes in more complex organisms, thereby contributing profoundly to our understanding of biological systems and phenomena.

Once a desirable mutant phenotype is identified through chemical mutagenesis, a key challenge lies in determining which among potentially thousands of induced mutations is responsible for the observed phenotype. (10) Identifying the causal mutation necessitates a substantial investment of labor and time, often demanding rigorous mapping and validation work to confirm the genetic change responsible for the observed phenotype (11, 12), (13). These limitations, especially when contrasted with RNAi (14, 15) or CRISPR-Cas9-based reverse genetics (16) methodologies, have led to the gradual obsolescence of EMS chemical mutagenesis and screens in the model organisms. RNAi, with its ability to knock down genes transiently and its amenability to high-throughput formats (17), enables the systematic analysis of gene function across the genome, offering a powerful tool for functional genomics (18). With CRISPR-Cas9, not only can specific genes be targeted, but specific types of mutations (e.g., point mutations, +deletions, insertions) can be introduced (19), offering a level of precision and control that is simply unattainable with chemical mutagenesis (20, 21).

On the other hand, chemical mutagenesis has unique advantages in the generation of a broad spectrum of alleles (22), including hypomorphic, hypermorphic, and neomorphic alleles, providing a rich reservoir for investigating gene function in a depth unattainable through complete gene knockdown or knockout strategies like RNAi or CRISPR-Cas9. Thus, forward genetic screens facilitate the exploration of nuanced gene functions (23), interactions (24) and pathway dynamics (25), often unveiling surprising connections and uncharted biological landscapes that are not immediately evident through targeted gene perturbation approaches (26). Furthermore, this strategy can illuminate the functionality of genes of unknown function or open reading frames that might be overlooked with hypothesis-driven reverse genetics (27). This stochastic, phenotype- driven approach allows for the serendipitous discovery of novel genetic interactions and pathways without preconceived notions about gene function, offering the potential to uncover entirely new facets of biology.

Whole-genome sequencing (WGS) has emerged as a formidable instrument for pinpointing causal mutations derived from genetic screens (28). In the context of *Caenorhabditis elegans* (*C. elegans*), several strategies have been devised to reduce the number of candidate causal mutations. One prevalent method involves mating mutants, which are in the N2 background, with the polymorphic Hawaiian strain (CB4856), while alternative strategies exploit DNA variants inherent to the initial background or introduced through mutagenesis (29). For the identification of suppressor mutations, the Sibling Subtraction Method (SSM) has been developed, which, by excluding genetic variants present in both mutants and their non-mutant siblings, markedly diminishes the roster of candidates (10). However, these strategies, given their reliance on genetic crosses, do not lend themselves to high-throughput applications. Given that the recent advancement of AlphaMissense has predicted millions of pathogenic variants (30), the incorporation of homologous ones into model organisms for genetic suppressor studies may enhance our comprehension of rescue mechanisms and forge paths towards the proposal of innovative therapeutic strategies. Consequently, there is a pressing demand for the development of scalable methods for identifying causal mutations.

In this study, we systematically analyzed *C. elegans* WGS data derived from 737 EMS- mutagenized worms, furnished by the Million Mutation Project (MMP) (22), alongside chromatin immunoprecipitation sequencing (ChIP-seq) datasets of DNA-binding proteins from the modENCODE project (31). Contrary to expectations, we discovered that EMS-induced mutations were not uniformly distributed across the *C. elegans* genome. Instead, a genome-wide EMS mutagenesis bias was uncovered, which correlates with both adjacent sequence context and chromatin structure, facilitating the development of a machine learning based model for predicting EMS-induced mutagenesis probabilities. This model enabled the generation of a genome-wide EMS mutagenesis probability map, which predicts the expected frequency of random mutations for each nucleotide within the *C. elegans* genome. Consequently, our method enhances our capability to discern causative mutations—enriched through genetic screening for a specific phenotype—from random mutations. By employing this pipeline to analyze WGS data of genetic suppressors that rescue a ciliary kinesin harboring a missense mutation in in *C. elegans*, we identified the causal mutations without the requisite establishment of recombinant inbred lines, thus enhancing the efficiency of high-throughput forward genetics.

## Result

### A non-uniform pattern of EMS-induced mutations across the *C. elegans* genome

We examined the distribution of a cumulative 265,169 Single Nucleotide Variants (SNVs) and insertion/deletions (InDels) across the 737 EMS-mutagenized genomes from the Million Mutation Project (Fig. 1A). A conspicuous non-uniform pattern emerged, characterized by ‘hot spots’—regions exhibiting a higher-than-average SNV density—and ‘cold spots,’ or regions manifesting a lower density. Given that EMS predominantly introduces an alkyl group at N7-guanine and O6-guanine, consequently inducing transitions from ‘G/C’ to ‘A/T,’ we postulated that mutation frequency might be modulated by the genomic concentration of ‘G/C’ nucleotides. Nonetheless, our correlation analysis revealed that the ‘G/C’ pair distribution only partially elucidated the mutation frequency (Fig. 1A, B). Moreover, we discerned variation in mutation frequency amongst different chromosomes (Fig. 1C), with chromosome X exhibiting an elevated mutation frequency and chromosome IV manifesting a diminished frequency, neither of which could be attributed to the ‘G/C’ distribution. These observations parallel the findings from two additional WGS datasets procured from our recent Artificial Evolution Summer School, in which undergraduates perform forward genetic screens for animals exhibiting dumpy (Dpy) or uncoordinated movement (Unc) phenotypes (*SI Appendix*, Fig. S1A-F). These results imply that EMS-induced variations are not uniformly distributed across the genome, potentially due to disparate local chromosomal features.

**Figure 1.**
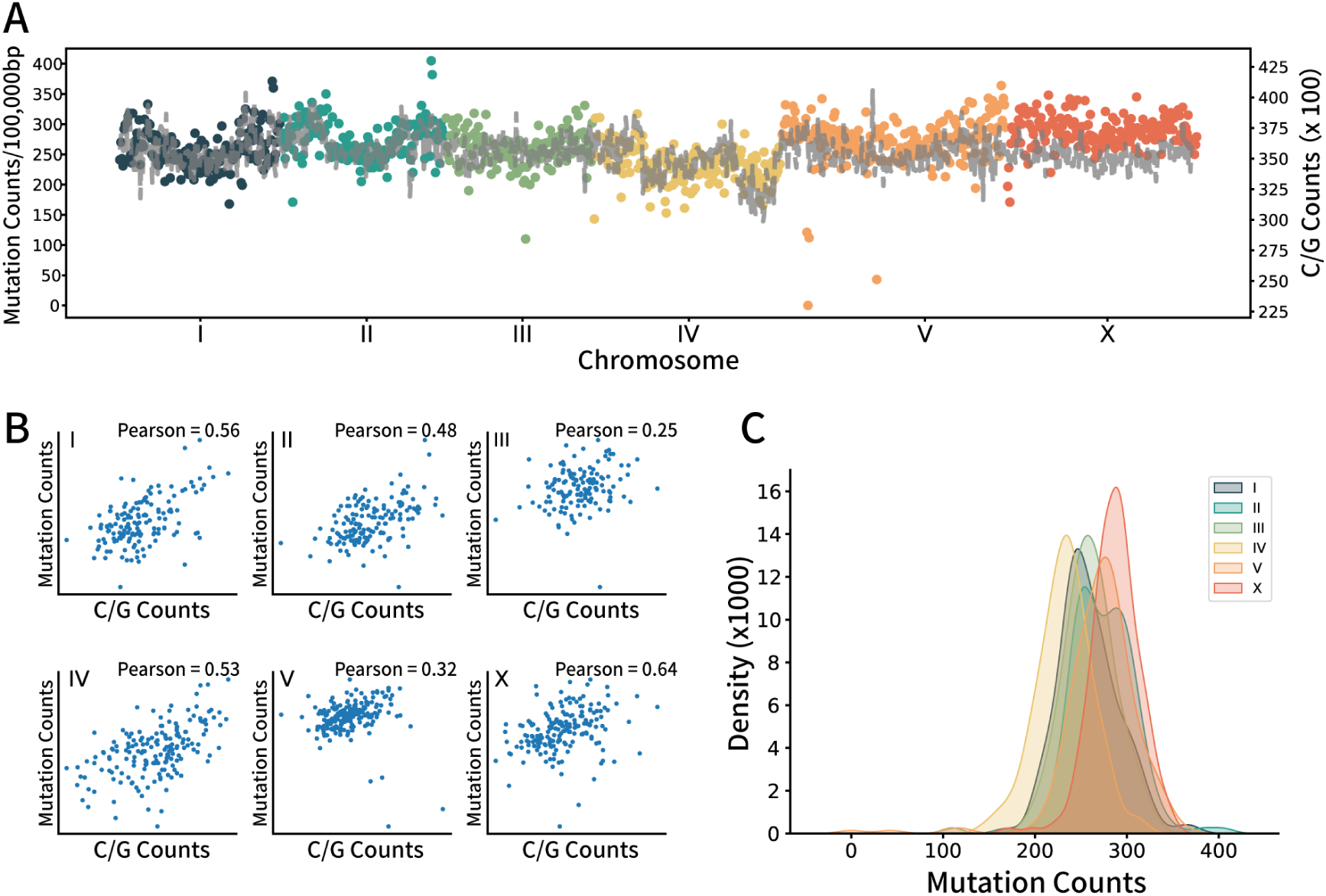
Uneven distribution of EMS-induced SNV. A) Scatter and line plot represents mutations in the MMP dataset. Scatter plot shows the mutation number on each chromosome. Each dot represents the number of mutated bases within every 100,000 bp counted from the first base pair of each chromosome. Line plot represents the number of ‘C/G’ base pairs in each 100,000 bp regions. B) Scatter plot represents the relationship between ‘C/G’ base pair contents and the number of mutations from the MMP data of each chromosome. Pearson correlation was used to evaluate the association between them. C) Kernel density estimation plot of the number of mutations on each chromosome.

### The adjacent sequence context influence the probability of EMS mutagenesis

We investigated the distribution of the four nucleotides both upstream (3’ +4/+3/+2/+1) and downstream (5’ -4/-3/-2/-1) of mutated bases across the genome (Fig. 2A, B). Should EMS-induced mutations transpire indiscriminately, uninfluenced by adjacent bases, an asymmetry-free distribution would be anticipated. However, our chi-square test illuminated that positions (+2/+1) and (-1/-2) adjacent to the ‘C/G’ nucleotide wield a significant impact on the efficacy of EMS mutagenesis. Examining the MMP dataset, we discerned that certain 5-base sequences exhibit a markedly heightened susceptibility to mutation. For example, sequences such as ‘AGGGG’ and ‘CCCCT’ are roughly tenfold more predisposed to mutation relative to sequences like ‘GTCGA’ and ‘TCGAC’ (Fig. 2C). The substantive impact on mutagenesis probability is not confined to the immediate neighbors of nucleotides. Even while maintaining the +1/0/-1 position constant, the +2 and -2 positions continue to sway mutagenesis probability (Fig. 2D, E), suggesting a viable avenue for predicting biases in mutation frequency.

**Figure 2.**
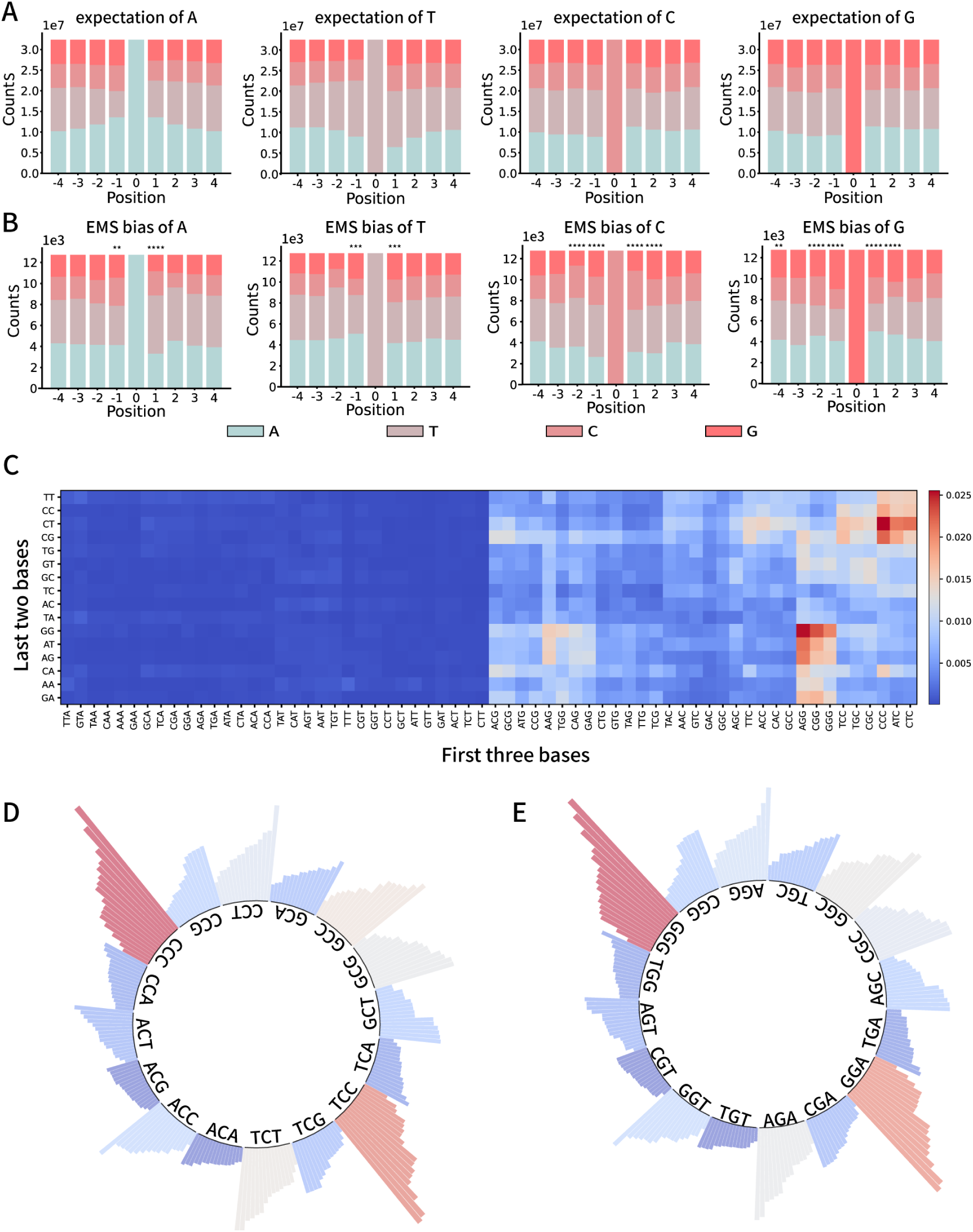
Flanking sequences affect EMS mutagenesis efficiency. A) The distribution patterns of flanking sequences of each kind of base (A, T, C, G) on the genome DNA. The number with ‘+’ or ‘-’ represents the base left (5’)/right (3’) to the position 0. These distributions were set as expectation values that if the mutagen select randomly on genome, it will cause mutations with the same distribution in flanking sequences. B) The distribution patterns of flanking sequences of each kind of mutated base in MMP dataset. Note that the distribution pattern is different from the corresponding distribution in (A). Chi-square test: * p<0.05, ** p<0.01, ***p<0.001, ****p<0.000. C) Heatmap recording the mutation rate of each 5-base pattern. D) Bar plots shows the variation of C mutagenesis rate when -1 and +1 position is fixed, as shown around the inner circle. E) Bar plots shows the variation of G mutagenesis rate when -1 and +1 position is fixed, as shown around the inner circle.

### Multiple DNA-binding protein patterns show correlation with the mutation events

Employing the adjacent sequence context, we crafted a graphical representation that elucidates the frequency of EMS-induced mutations. To exemplify the methodology, we showcase the map in relation to mutation events across a broad range on chromosome V, spanning from position 1,800,000 to 3,450,000 (Fig. 3A, B). We incorporated a representative gene, *ttn-1*, situated on chromosome V between positions 6,120,909 and 6,202,632 (*SI Appendix,* Fig. S2A, B). In both scenarios, the maps reveal expansive genomic segments characterized by a notable diminution in EMS-induced mutations, the ‘silent region’ of which belies the predictions formulated on adjacent sequence context.

**Figure 3.**
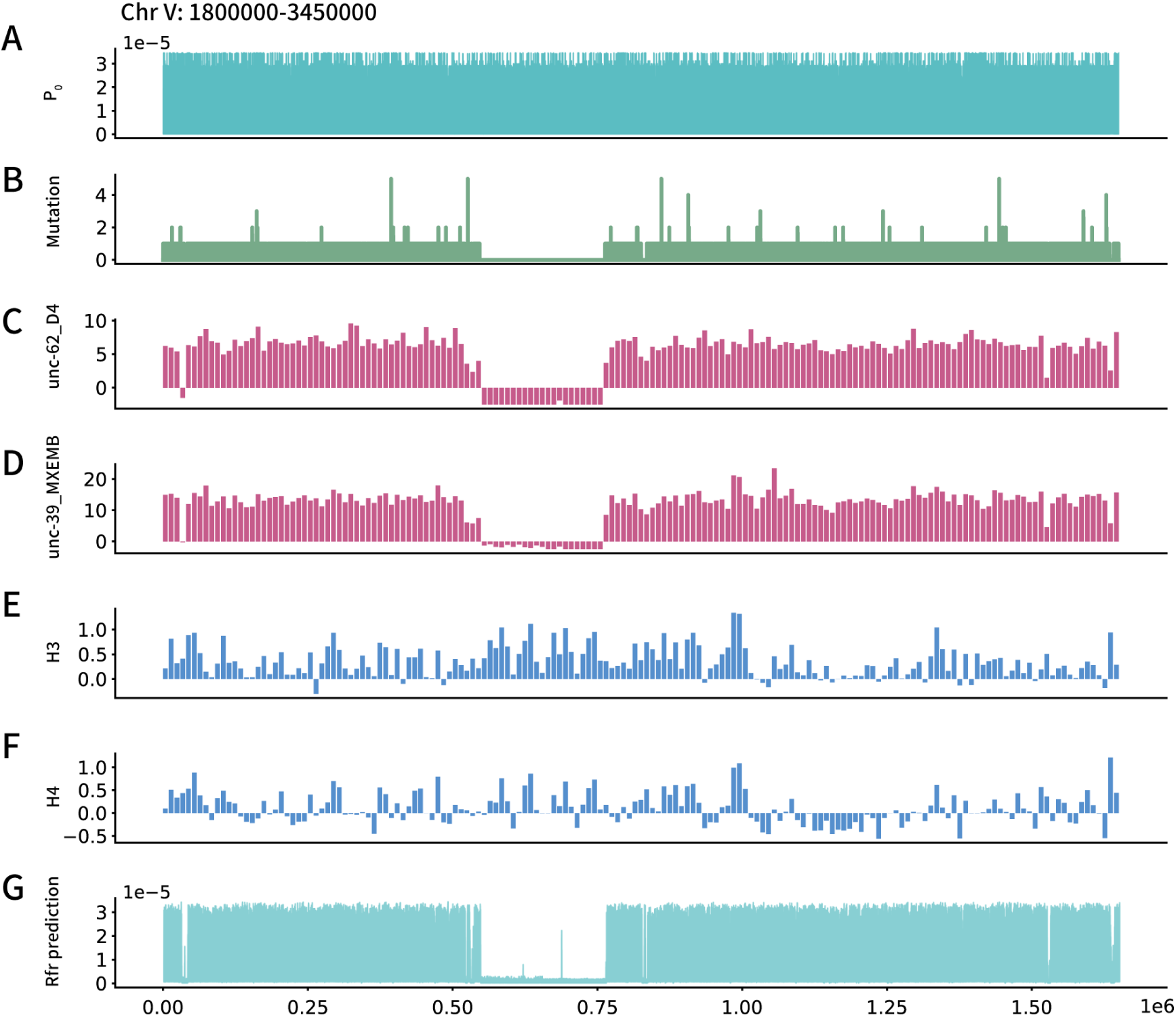
Mutation numbers in the MMP data shown association with the DNA-binding protein features. Take the MMP data and DNA-binding protein features on Chr V:1800000∼3450000 as an example A) Line plots shows the prediction made from the Flanking sequence preferences of EMS mutagenesis (*P*_0_). B) The mutations observed in the MMP dataset. C) Raw CHIP-chip signal data of unc-62 binding in young adult worms. D) Raw CHIP-chip signal data of unc-39 binding in worm embryo. E) Raw CHIP-seq signal data of histone H3 in L3 larval worms. F) Raw CHIP-seq signal data of histone H4 in L3 larval worms. G) Line plots shows the prediction made by a Random Forest regressor trained by DNA-binding protein data.

We posited that a unique chromatin state might elucidate the emergence of these ‘silent regions’. Tightly supercoiled DNA, potentially having circumscribed interactions with extraneous substances (32), along with particular DNA-binding proteins, might confer protection to their target sequences, rendering them less vulnerable to chemicals that engage with DNA (33). Consequently, we probed the *C. elegans* CHIP-seq datasets of DNA-binding proteins and histone modifications from the modENCODE project (Fig. 3C-F and *SI Appendix* , Fig. S2C-F). On chromosome V, a silent region extends over 200kb in length (Fig. 3B). Within this expanse, the binding activity of the transcription factor UNC-62 and UNC-39 manifests irregularly, as evidenced by the CHIP-seq signal (Fig. 3C, D). The absence of CHIP-seq signals in this area might denote a distinctive chromatin state that is less amenable to external substances. Analogously, the ‘silent region’ of the gene ttn-1 also undergoes a comparable albeit less marked depletion of transcription factor binding (*SI Appendix*, Fig. S2C, D). Nonetheless, a distinctive histone binding pattern is discernible in this area (*SI Appendix,* Fig. S2E, F), a pattern not observed in the preceding 200kb ’silent region’. These singular DNA-binding protein features might suggest that an alternative chromatin state might contribute to a low mutagenesis frequency in a certain region. Probing mutations and the attributes of DNA-binding proteins suggests that regions with elevated or reduced mutation probabilities may originate from direct causative factors or mere coincidence. Nevertheless, it seems judicious to elucidate the binding patterns of these proteins and explore their association with mutation probability.

Using CHIP-seq dataset collections from the Wormbase, we next examined the potential correlation between DNA-binding protein features and mutagenesis probability with machine learning techniques.

Nevertheless, it seems judicious to elucidate the binding patterns of these proteins and explore their association with mutation probability. Employing CHIP-seq dataset compilations from Wormbase, we subsequently scrutinized the potential correlation between DNA-binding protein characteristics and mutagenesis probability through the lens of machine learning methodologies.

### Random Forest regressor (Rfr) modeling to predict the mutation frequency

Employing CHIP-seq dataset compilations from modENCODE and Wormbase (31), we subsequently scrutinized the potential correlation between DNA-binding protein characteristics and mutagenesis probability through the lens of machine learning methodologies. We developed a regression model using the Random Forest Regressor, a machine learning algorithm that employs ensemble learning techniques for predictive tasks (34). The underlying hypothesis is that if a group of nucleotides (in this case, group size is ≥ 30,000) exhibit identical properties and coexist in the same environment context, they will exhibit an equal susceptibility to EMS-induced mutations. The Random Forest model stratifies data according to specific conditions and computes the average mutation probability for each one (Fig. 4A, Materials and Methods).

**Figure 4.**
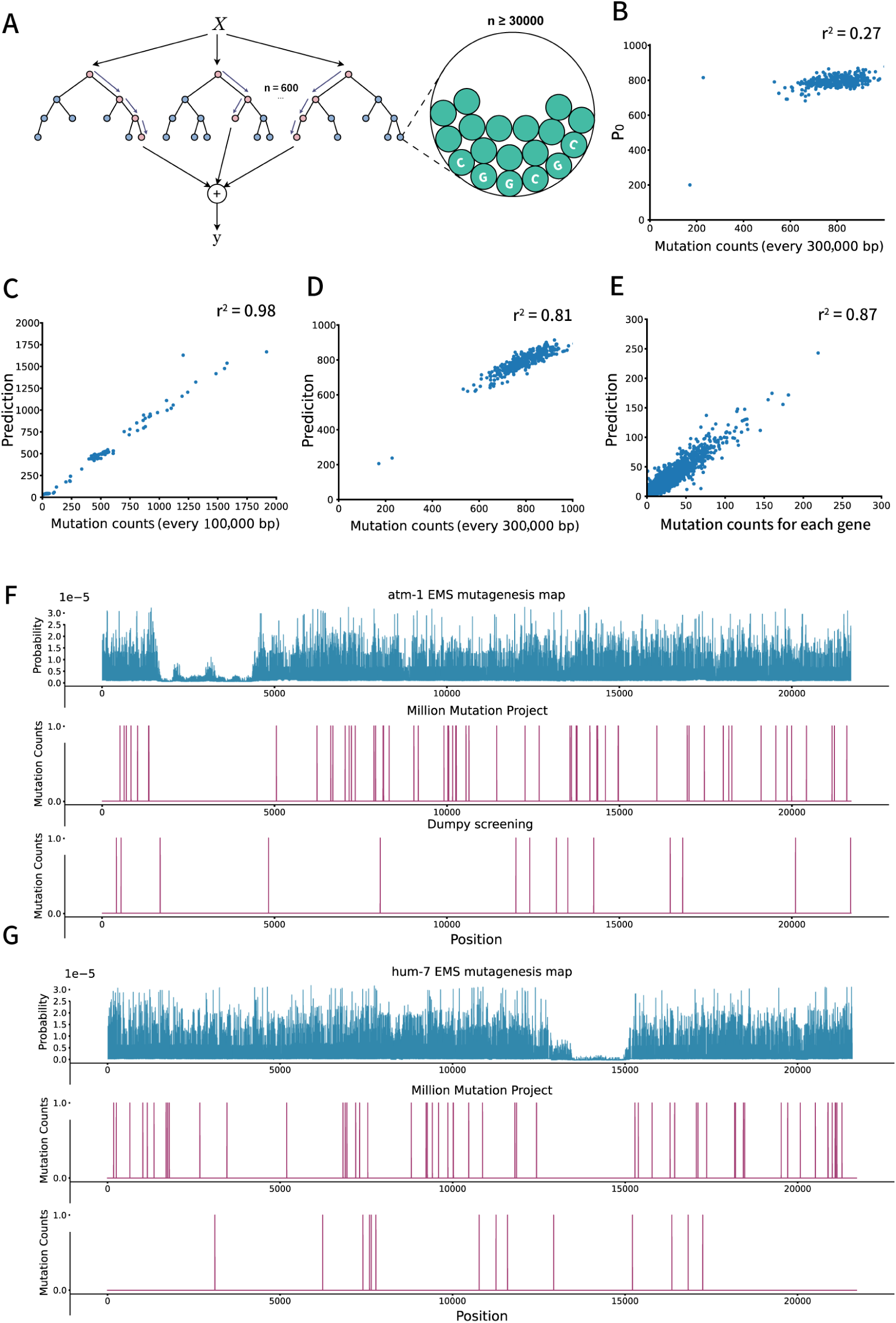
Random Forest modeling and validation. A) Schematic diagram of the Random Forest regressor (RFr) model. 600 trees were modeled and each tree randomly used at most 70% percent of the 24 features (≤16 features). The miniest leaf size is set to be 30000 to avoid overfitting. In this model, each decision tree in the forest will put bases with similar properties (flanking sequences and DNA-binding protein patterns) together into a output node, and use the average mutation rate in each node as output. Consequently, the forest will take the average output of every 600 trees as the final output, with which the mutation rate of this kind of base can be predicted. B) Prediction made only by *P*_0_. The worm genome was divided into 300,000 bp blocks and each dot represents the expectation of mutation number made by *P*_0_ and the actual mutation counts in MMP dataset. C) The performance of RFr on test set, which contains 20% of the whole dataset. The test set was sorted by the prediction and was divided into 100,000 bp blocks. Each dot represents the expectation of mutation number made by RFr and the actual mutations counts in MMP dataset in each 100,000 bp long blocks. D) Scatter plot of the overall performance of RFr on the whole genome. The genome was divided into 300,000 bp blocks. Each dot represents the expectation of mutation number made by RFr and the actual mutations counts in MMP dataset in each 300,000 bp long blocks. E) Scatter plot of the overall performance of RFr on 19639 genes of the *C. elegans*. Each dot represents the expectation of mutation number made by RFr and the actual mutations counts in MMP dataset of each gene. F) Representative EMS mutagenesis map of gene *atm-1*. Followed by the mutation map in the MMP dataset and dumpy screening. G) Representative EMS mutagenesis map of gene *hum-7*. Followed by the mutation map in the MMP dataset and dumpy screening.

As depicted in the schematic diagram (Fig. 4A), nucleotides with analogous flanking sequences and DNA-binding protein attributes are categorized into the same output node of each tree. The computed mean mutagenesis frequency for that node subsequently serves as the prognosticated mutagenesis frequency for any nucleotide with similar characteristics that are allocated to the same node during ensuing predictions. To formulate our model, we leveraged a dataset, encompassing 24 DNA-binding protein datasets from modENCODE, which includes information about histone binding, modification, transcription factor interactions, and additional epigenetic modifications (*SI Appendix,* Table S1). We constructed 600 decision trees, each predicated on the aforementioned methodology, with each tree being trained on a randomly chosen subset of up to 16 protein features.

The ultimate model output can be succinctly represented as 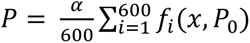 (see also Materials and Methods), which involves averaging the predictions from each individual tree to provide a more precise overall prediction. In this equation, the variable α factor in experimental variations stemming from fluctuations in EMS concentrations, developmental stages, ambient temperatures, and other potential elements that may impact EMS mutagenesis effectiveness.

To validate the model’s performance, our initial validation assessed predictions grounded solely on *P*_0_, demonstrating congruent patterns across every 300 kb genomic block. These patterns were juxtaposed with the discerned fluctuations in mutation occurrences within the MMP dataset (Fig. 4B). Upon training utilizing a randomly selected 80% of the data, the model’s performance was appraised against the test set. The test set was categorized based on prediction results and subsequently partitioned into 100kb blocks. Anticipated mutation tallies were then contrasted against actual mutation instances documented in the MMP dataset (Fig. 4C). Further, a thorough evaluation juxtaposed the anticipated mutation tallies within 300kb genomic blocks (Fig. 4D) and individual genes, as per the WBcel235 annotation (Fig. 4E), against actual mutation tallies from the MMP dataset. Collectively, these comparisons refine the rudimentary P0 map, facilitating a meticulous prediction of EMS mutagenesis probability at individual nucleotides (Fig. 3G, *SI Appendix,* Fig. S2G, Fig. 4 F, G).

Subsequently, we selected two representative genes and created EMS mutagenesis maps, highlighting variations in mutagenesis frequency across the entire gene. These maps were then compared to mutations observed in the MMP dataset and sequencing data from the dumpy screening mentioned earlier (Fig. 4F, G). In regions identified as ‘silent region’ based on our predictions, neither screening approach revealed mutations. Conversely, due to the limited number of SNVs in the MMP dataset, numerous nucleotides remained unmutated in this dataset. Notably, the continuous absence of observed mutations on the actual mutation occurrence map did not diminish the model’s ability to predict the potential for mutagenesis: In the dumpy screening, we observed mutations in regions where the MMP dataset displayed minimal mutations (Fig. 4F, G). This suggests that our model recognized the specific characteristics of nucleotides in these regions and transferred the knowledge that similar nucleotides were prone to mutation in other genomic regions in order to make accurate predictions.

### The EMS mutagenesis probability map facilitates the identification of causal mutations

We employed the EMS mutagenesis probability map to discern the causal mutation within a genetic suppressor screen, amending defects instigated by a missense mutation in the ciliary kinesin OSM-3. This kinesin drives intraflagellar transport, a process pivotal for the construction of olfactory cilia in *C. elegans* sensory neurons. These segments harbor an abundance of G protein-coupled receptors (35), enabling the organism to perceive environmental stimuli, inclusive of various odorant molecules. A functional impairment of the OSM-3 kinesin culminates in a specific loss of distal ciliary segments (36); concomitantly, the delocalization of GPCRs or the prevention of odorant-receptor interaction causes animal behavioral defects, such as an incapacitation in executing osmotic avoidance (Osm).

Our preceding genetic screens isolated the E251K missense mutation within the motor domain of OSM-3 kinesin. This mutation parallels a pathogenic variant, E253K, identified in the KIF1A kinesin (37), mutations within which give rise to a spectrum of neurological disorders collectively recognized as KIF1A-Associated Neurological Disorder (KAND). The E253K mutation in KIF1A is hypothesized to destabilize the structure of the back door, thereby suppressing γ-phosphate release and subsequent ATP binding within the motor domain (38, 39). An in vitro single-molecule motility assay revealed that E253K induces a stringent binding to the microtubule (MT) yet precludes the engagement in processive motion of KIF1A. Consistently, *C. elegans* harboring the E251K mutation in OSM-3 disrupts their distal ciliary segments with full penetrance. Whereas wild-type animals exploit their distal ciliary segments to uptake the fluorescent dye DiI from the culture medium (40), all examined OSM-3 E251K mutant animals failed to do so (N > 500), manifesting a Dyf phenotype detectable under a fluorescence stereoscope.

Utilizing the Dyf phenotype as an efficacious readout probing ciliary defects, we executed genetic suppressor screens aimed at restoring dye-filling capacity and recuperating ciliary distal segments. Our screens isolated 38 independent suppressors. We conducted whole-genome sequencing of all suppressors and employed the EMS mutagenesis probability map to analyze the WGS data. Initially, background mutations, defined as those shared across a wide range of samples, were identified and removed. Subsequently, we evaluated the effectiveness of EMS mutagenesis (Materials and Methods). The remaining mutations were compared to the adjusted mutagenesis map (Fig. 5A). To assess the enrichment effect on each gene by the genetic screening, we calculated the fold change and p-value for each gene showing mutations in the background-removed mutation pool (Fig. 5A).

**Figure 5.**
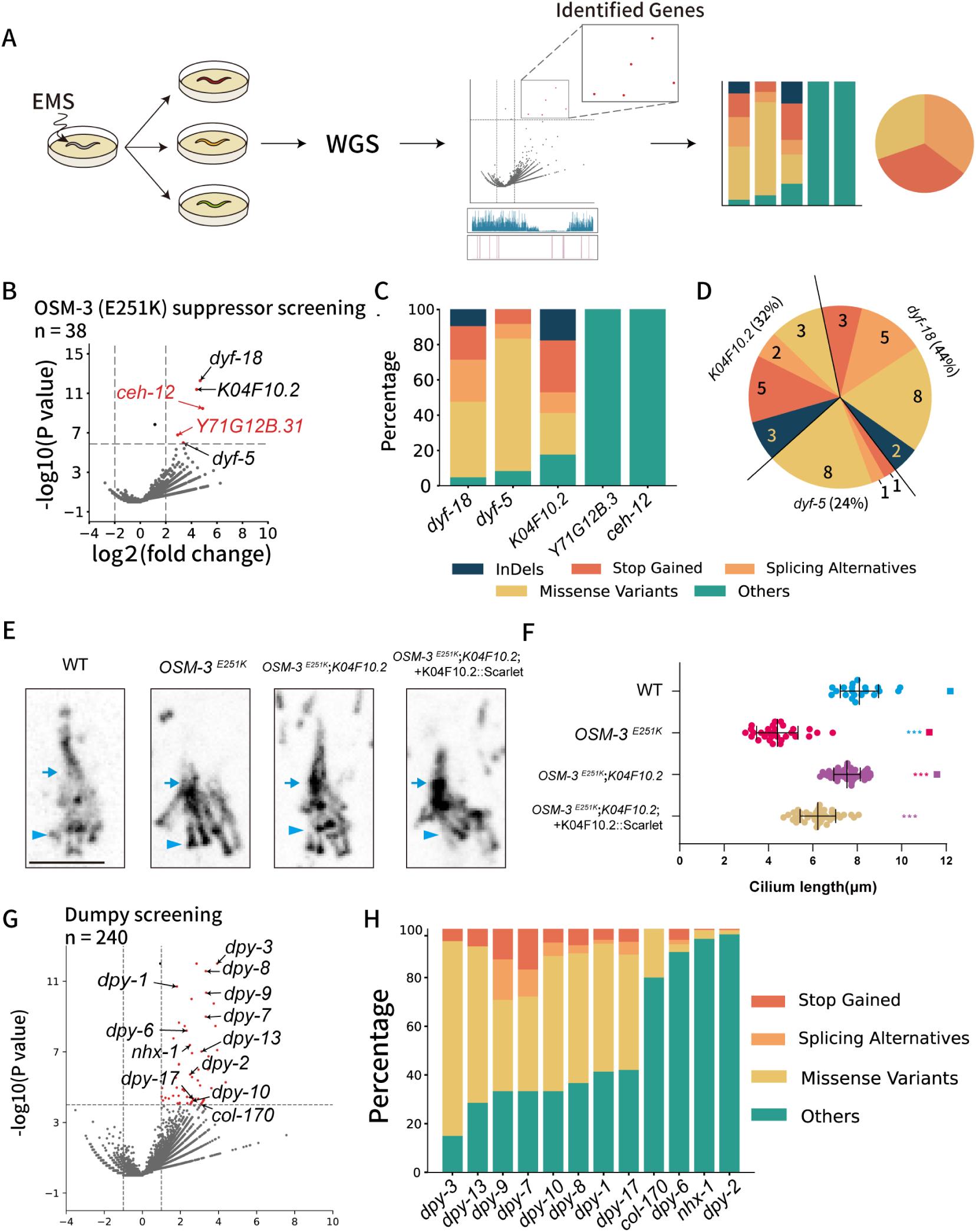
Predict target genes with the EMS mutagenesis map. A) Schematic diagram of the analysis pipeline. After genetic screening of the EMS mutated worms, those with a target phenotype were sequenced and mutation information were pooled. Then it can be compared with the EMS mutagenesis map based on the RFr (see Materials and Methods). Finally, candidate genes’ mutation profile was analyzed by the loss of function impact of InDel, stop-gain, splicing, missense mutation. B) Volcano plot showing the result of an OSM-3 (E251K, cas22599, n=38) suppressor screening comparing to the EMS mutagenesis map. There were 5 genes showing significant difference between mutation expectation and actual mutation number. C) Bar plot showing the mutation properties of these five genes. D) Taking all the mutations inducing a high impact on protein into consideration, each of the 38 samples have one high-impact mutation in *dyf-5, dyf-18* and K04F10.2. E) Ciliary defects in the OSM-3 (E251K, cas22599) mutant animals rescued by a K04F10.2 mutated allele (Arg506*, cas23441). K04F10.2 gDNA overexpression in cas23441 exhibits shorter cilia. Arrowheads indicate the ciliary base and transition zone, and arrows indicate junctions between the middle and distal segments. F) Cilium length (mean ± SD) in each group, n = 30 to 50. *** P<0.001 one-way ANOVA. G) Volcano plot showing the result of a dumpy screening (N=240) from EMS mutated N2 wildtype strain comparing to the EMS mutagenesis map. Dumpy-related or body wall related genes were labeled. H) Bar plot showing the mutation properties of genes labeled in (G).

Our analysis unveiled the top five candidate genes, which demonstrated a markedly biased EMS mutagenesis rates in our suppressor screen but did not exhibit any enriched mutation frequency in our Dpy or Unc screens (Fig. 5B). Among these candidates, two ciliary kinases, DYF-5 and DYF-18, have been recognized for their efficacy in rescuing ciliary defects in *osm-3* mutant animals. We obtained 10 and 16 mutant alleles for dyf- 5 and dyf-18, respectively. The majority of these mutant alleles induce missense mutations, introduce stop codons, or cause splicing mutations within the coding region. These results underscore that the EMS mutagenesis probability map facilitates the identification of causal mutations in genes known to participate in this process. The remaining three genes are not well characterized, and it is unclear which one or ones might be implicated in the regulatory mechanisms of OSM-3.

Through categorizing the mutational characteristics of three genes, we observed that two of them harbored mutations - other than missense, introduced stop, or splicing mutations - which are unlikely to disrupt the function of the gene products. In stark contrast, 12 genetic suppressors encompass various loss-of-function mutations within the coding region of the K04F10.2 gene. Noteworthily, given the substantial size of the initial group subjected to screening, the concurrence of two suppressor alleles within the same strain emerges as exceptionally rare. Notably, the mutations with a significant impact did not coincide across the 38 suppressor strains (Fig. 5D), suggesting that the 38 genetic suppressors induce defects in three genes: dyf-5, dyf-18, and K04F10.2. Given that we have already procured 12 distinct alleles of K04F10.2, and generating additional mutants of this gene may not yield further insight, we endeavored to conduct transgenic rescue experiments to further determine whether K04F10.2 operates as the suppressor gene. To this end, we introduced the genomic DNA of K04F10.2, tagged with the red fluorescent protein Scarlet and controlled by its 2kb endogenous promoter, into OSM-3 E251K; K04F10.2 double mutant animals (Fig. 5E, F). The IFT-dynein heavy chain CHE-3, which traverses the entire length of cilia, was marked with green fluorescence and used as a ciliary marker to ascertain ciliary length (Fig. 2E) (41). As anticipated, GFP fluorescence was observed along cilia measuring 8.09±0.86 μm. In mutants harboring the OSM-3 E251K mutation, the distal ciliary segment was absent, resulting in cilia of the shorter length (4.40±0.93 μm). Interestingly, the introduction of wild-type K04F10.2 into the OSM-3 E251K; K04F10.2 double mutants restored ciliary length to 7.54±0.60 μm. However, the ciliary length was reduced to 6.22±0.81 μm, similar to OSM-3 E251K single mutants (Fig. 2F), suggesting that K04F10.2 mutations act as suppressors for OSM-3 E251K. These results demonstrate that the EMS mutagenesis probability map is instrumental in isolating causal mutations in genes previously unlinked to specific processes.

## Discussion

In conclusion, this research delineates the non-uniform mutation rate instigated by EMS mutagenesis across the *C. elegans* genome, underscoring that both proximate sequence context and chromatin status are ostensibly correlated with mutation frequency. Leveraging these factors, we deployed a Machine Learning assisted pipeline to sculpt a genome-wide EMS mutagenesis probability map. Operating under the premise that causative mutations will achieve enrichment through genetic screens pertinent to a specific phenotype amidst random mutations, we utilized the map to scrutinize Whole Genome Sequencing (WGS) data derived from *C. elegans* forward genetic screens. The findings illuminate that the map expediently facilitates the discernment of genes known for their instrumental roles in these processes. Notably, the map also prognosticated a novel gene entwined in a given regulation, a prediction substantiated through transgenic rescue experiments. This holistic approach to causal gene identification eschews labor-intensive genetic crossing and demonstrates that bioinformatic analysis of WGS data via the EMS mutagenesis probability map can present a potent conduit for directly interfacing phenotype with genotype. This enables scalable causal gene identification, thereby reinvigorating the utility of forward genetic screens.

The elucidation of K04F10.2 as a suppressor gene mitigating ciliary defects induced by the OSM-3 E251K mutation inaugurates avenues for mechanistic investigations. Previous research has illuminated that the abrogation of ciliary kinases DYF-5 or DYF- 18 permits an alternative ciliary kinesin-II to ectopically enter the ciliary distal region (42), thereby substituting the functionality absent in OSM-3. In essence, these two ciliary kinases may not exert their influence directly upon OSM-3. Rather, they curtail a concurrent ciliary transport pathway, and their loss enables kinesin-II to compensate for the OSM-3 deficit. Consequently, in the singular mutants of dyf-5 or dyf-18, the organisms exhibit aberrantly elongated cilia (42). In contrast, K04F10.2 might orchestrate its function through a mechanism disparate from these two ciliary kinases. We and others have not been able to discern any overt ciliary defects in the single mutant defective in K04F10.2. A pioneering systematic exploration of ciliary genes has annotated K04F10.2 as a conserved putative binding associate of a microtubule-severing protein, namely katanin, hinting at a role in microtubule regulation (43). Nonetheless, the underlying mechanisms remain mysterious. Considering that the E251K mutation in OSM-3 is analogous to the E253K mutation in KIF1A, future studies will ascertain whether the inhibition of K04F10.2 can ameliorate neuronal defects induced by KIF1A E253K, potentially unveiling a novel therapeutic target for intervening in KIF1A E253K-associated neurological disorders.

Recent endeavors to conduct genetic suppressor screens for various missense mutations within OSM-3 have been undertaken, inclusive of published suppressors of a motor hinge mutation, OSM-3 G444E (44, 45). Nonetheless, no mutations of K04F10.2 were unveiled as suppressors for the ciliary defects attributed to OSM-3 G444E, positing that K04F10.2 might exert its suppressive effects on the E251K mutation in OSM-3 in a residue-specific modality. This observation accentuates the imperative of functional residuomics: each residue may exhibit intrinsic uniqueness, and mutations at each individual residue may be ameliorated via divergent, distinct mechanisms.

Given the ubiquitous chemical principle underpinning EMS’s capacity to induce genetic mutations across diverse species, it is conceivable that the adjacent sequence context and chromatin status might also engender a non-uniform distribution of EMS- induced alterations throughout all genomes. Congruent with our findings, the efficacy of EMS mutagenesis in rice (*Huanghuazhan*) is correlated with its flanking sequence and chromatin status (46). Consequently, it may not be startling to observe that systematic analyses of published whole-genome sequencing (WGS) data across species divulge a pervasive bias in EMS mutation rates at genome-wide levels. Identifying EMS-induced causal mutations in species characterized by complexities in genome size, ploidy number, and life cycle surpassing those of *C. elegans* poses a substantially more formidable challenge. Therefore, formulating analogous EMS mutagenesis probability maps for these species could markedly expedite forward genetics endeavors therein.

## Materials and Methods

### WGS data from the Million Mutation project

WGS data from the Million Mutation project can be accessed via the official webpage of Simon Fraser University (http://genome.sfu.ca/mmp/). For our study, we utilized mutation data from a total of 737 strains isolated after EMS mutagenesis was used to train and test the model. Raw sequencing data in fastq format are available from Home - SRA - NCBI (nih.gov). The average sequencing depth of these data were about 15x fold and 265169 variations were found in this EMS mutation dataset.

### DNA-binding protein data

*C. elegans* CHIP-chip and CHIP-seq data can be obtained from the modENCODE project (www.modencode.org) and visualized in Wormbase Genome Browser (JBrowse (wormbase.org)).

### *P*_0_ calculation from the MMP dataset

Flanking sequence preferences of EMS mutagenesis (referred to as *P*_0_) were calculated with the MMP dataset and *C. elegans* genome WBcel235 (*<u>Caenorhabditis_elegans</u>* <u>- Ensembl Genomes 57</u>) as

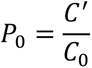

*C*’ represents the count of a specific five-base pattern underwent mutation in the MMP dataset and *C*_0_ is the number of the same pattern in the reference genome.

### Preparation of DNA-binding protein data and Random Forest Regressor modeling

The raw CHIP-chip and CHIP-seq data were preprocessed to better reflect the actual distribution of DNA-binding protein on the genome. A python script was employed to apply a moving average with the window size of 100 bp to smooth the data across each chromosome. The smoothed DNA-binding protein dataset was then utilized to train and validate a random forest regressor after randomly spilt the data into a 80% training set and a 20% test set.

The random forest comprised a total of 600 decision trees, each trained on the provided training dataset. To prevent overfitting, we implemented feature selection by randomly considering up to 70% of the available features for each tree. Additionally, to maintain model integrity, we set a minimum leaf size of 30,000 instances. To produce the final prediction, the output from each individual tree was aggregated. This ensemble approach allowed us to generate a comprehensive mutation probability assessment for each type of nucleotide.

During validation, the test set, comprising both mutation information and additional features, was sorted based on their predicted mutation rates. Subsequently, the test set was divided into 100kb blocks. To validate the model’s predictions, we computed the mutation expectation and the actual mutation count in the MMP dataset. By calculating the likelihood of mutation for each individual base pair, we were able to determine the mutation rate for each block, or for any genomic sequence length, using the formula:

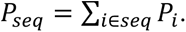

This approach allowed us to assess the model’s predictive accuracy across different genomic regions.

During a genetic screening involving the potential mutation of millions of base pairs, the number of variations for each nucleotide can be reasonably modeled to follow a binomial distribution. Consequently, the expected mutation count can be calculated as:

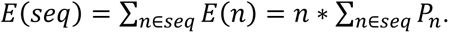

This formula is applicable within a population of n strains, and it allows us to estimate the anticipated number of mutations across the sequence under investigation.

The EMS effectiveness can be subject to variation due to factor such as changes in temperature, the age of the worms, minor temporal differences, and alterations in mutagen concentration during the EMS mutagenesis process. To assess this effectiveness, we compare the average number of mutation events per strain in different batches of EMS mutation experiments. Here we denote the mutagen effectiveness of the MMP strains as *α*_0_ and the efficiency in another experiment as *α*_1_. The random forest regression model takes the form of:

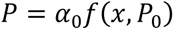

*x* represents the DNA-binding protein features used by the model to make predictions and adjust the initial prediction of *P*_0_. This model can also be applied to another dataset as:

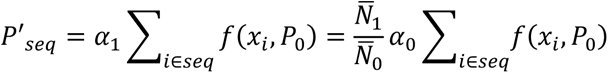

In this case, *N̄*_0_ and *N̄*_1_ represent the average number of mutation events that occurred in the MMP strains and the batch to be analyzed, respectively.

### Model evaluation and feature importance

To evaluate the accuracy of this regression model, we compare the expectation and the actual mutation count within different regions of the *C. elegans* genome, resulting in the determination of the coefficient of determination (*R*^2^), given by:

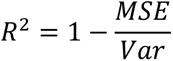

Additionally, we assess the importance of each feature used in making predictions through permutation scores. In this analysis, each feature is successively replaced with random noise sharing the same value distribution as the original data. The model, initially trained on the original dataset, is then employed to make predictions using the dataset in which a feature has been replaced with noise. The permutation importance of feature *j* is subsequently calculated as:

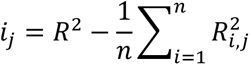

Where feature *j* undergoes *n* shuffling iterations.

Python scripts and the model are accessible on GihHub at (young55775/Genetorch-developing (github.com)). The mutation and feature importance data (*SI Appendix,* Fig. S3), have been normalized to a range of 0 to 1 using the formula:

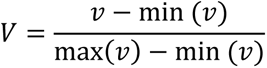

### Worm culture

*C. elegans* were maintained under a consistent temperature of 20 ℃ according to the standard method. Nematode growth medium (NGM) with *Escherichia Coli* OP50 seeded on it was used to cultivate these worms. *C. elegans* strains used in this study are listed in *SI Appendix,* Table S2.

### EMS mutagenesis

Worms synchronized at the late L4 stage were carefully collected with 4 mL M9 buffer (*SI Appendix,* Table S1). Subsequently, these collected worms were placed in 50 mM EMS buffer at room temperature with continuous rotation for a duration of 4 hours. Following this treatment, the worms underwent a thorough washing process with M9 buffer and were then cultured under standard conditions. Approximately 20 hours later, the adult worms were subjected to a bleaching procedure to isolate their eggs (referred to as F1). These eggs were subsequently distributed across approximately 100 separate 9 cm NGM plates, with an average of 50 to 100 eggs placed on each plate. Any adult worms displaying the desired phenotype were meticulously collected and placed in individual culture settings. After a careful examination of their offspring, these individuals were subjected to sequencing using an Illumina next-generation sequencer.

### WGS data analyze

Mutation data of MMP dataset were directly downloaded from the MMP homepage (http://genome.sfu.ca/mmp/mmp_mut_strains_data_Mar14.txt). Raw reads obtained from the next-generation WGS were assessed for duplication and quality with FastQC and were trimmed using Trim_galore (version 0.4.4) to remove the adaptor sequence and low-quality reads. After that, clean reads were aligned to the reference genome (WBcel235) using BWA-MEM2 (version 2.2) with default parameters. In all these sequencing results, > 20x average coverage is ensured. Variations were detected using freebayes (version 1.3.6) and annotated using SnpEff. Finally, a filter was set to ensure the variation quality and limit false positive rate: only sequence depth >5 and allele frequency > 0.8 variations were taken into further analyze.

### Strain construction

To perform the rescue experiment, we synthesized the pK04F10.2::K04F10.2::Scarlet DNA fragment using SOEing PCR (47) and subsequently created the transgenic strain through microinjection.

## Author contribution

G.O. and Z.G. designed the project. Z.G. and Y.W. analyzed sequencing data. Z.G. built the model. S.W. performed genetic screening. S.W. and Z.W. conducted the rescue experiment. G.O. , Z.G. and Y.W. wrote the paper.

## Acknowledgments

This work was supported by the National Natural Science Foundation of China Grants (31991190, 31730052, 31861143042, 31671444, and 31871352) and National Key R&D Program of China Grants (2019YFA0508401 and 2017YFA0102900).

## Declaration of Interests

The authors declare that they have no competing interests.

---

**Figure S1.**
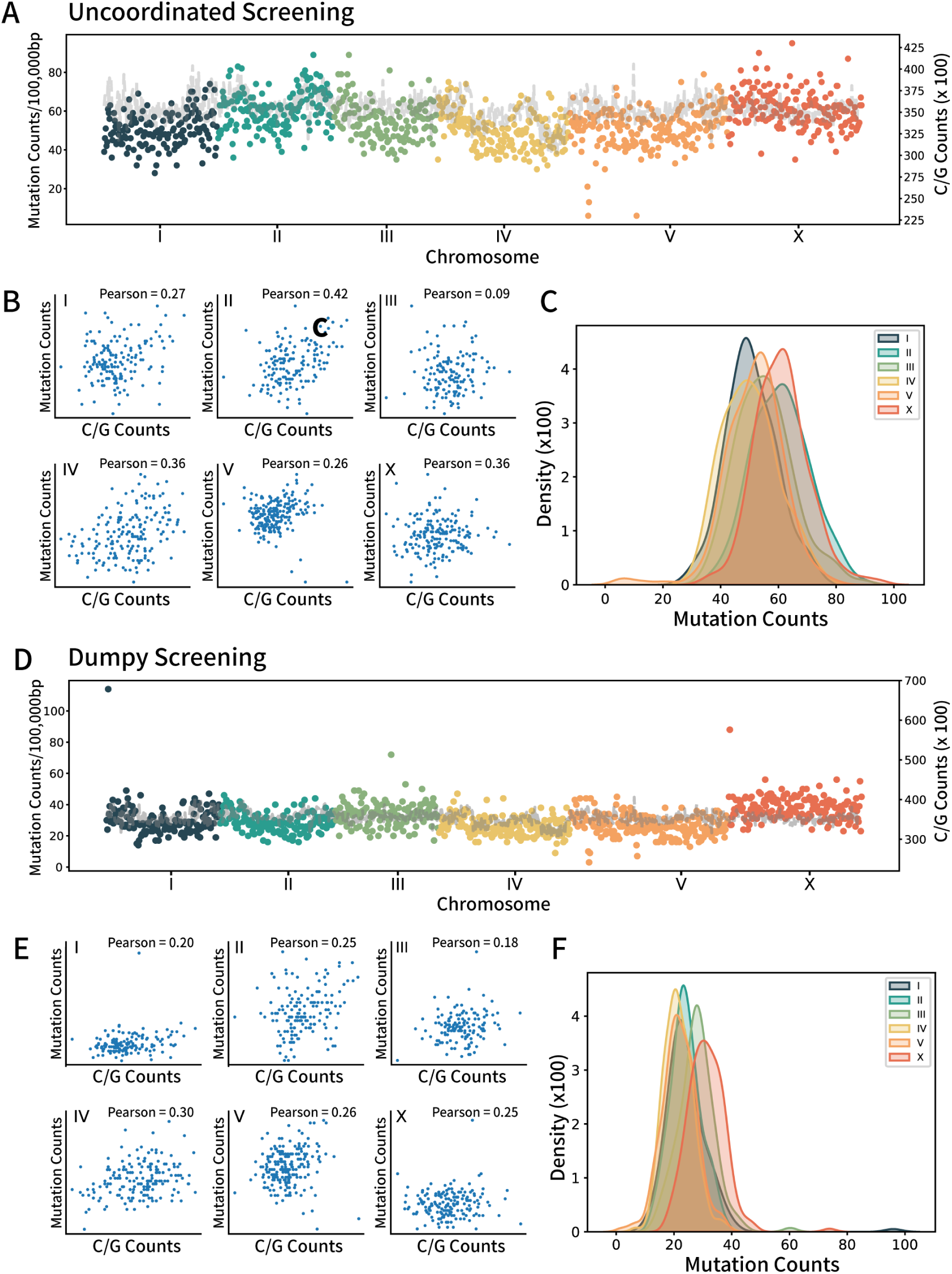
Uneven distribution of EMS-induced SNV in screening of dumpy and uncoordinated phenotype. A) Scatter and line plot represents mutations in the Uncoordinated screening dataset (n=118). Scatter plot shows the mutation number on each chromosome. Each dot represents the number of mutated bases within every 100,000 bp counted from the first base pair of each chromosome. Line plot represents the number of ‘C/G’ base pairs in each 100,000 bp regions. B) Scatter plot represents the relationship between ‘C/G’ base pair contents and the number of mutations from the Uncoordinated screening data of each chromosome. Pearson correlation was used to evaluate the association between them. C) Kernel density estimation plot of the number of mutations on each chromosome in uncoordinated screening dataset. D) Scatter and line plot represents mutations in the dumpy screening dataset (n=240). Scatter plot shows the mutation number on each chromosome. Each dot represents the number of mutated bases within every 100,000 bp counted from the first base pair of each chromosome. Line plot represents the number of ‘C/G’ base pairs in each 100,000 bp regions. E) Scatter plot represents the relationship between ‘C/G’ base pair contents and the number of mutations from the dumpy screening data of each chromosome. Pearson correlation was used to evaluate the association between them. F) Kernel density estimation plot of the number of mutations on each chromosome in dumpy screening dataset.

**Figure S2.**
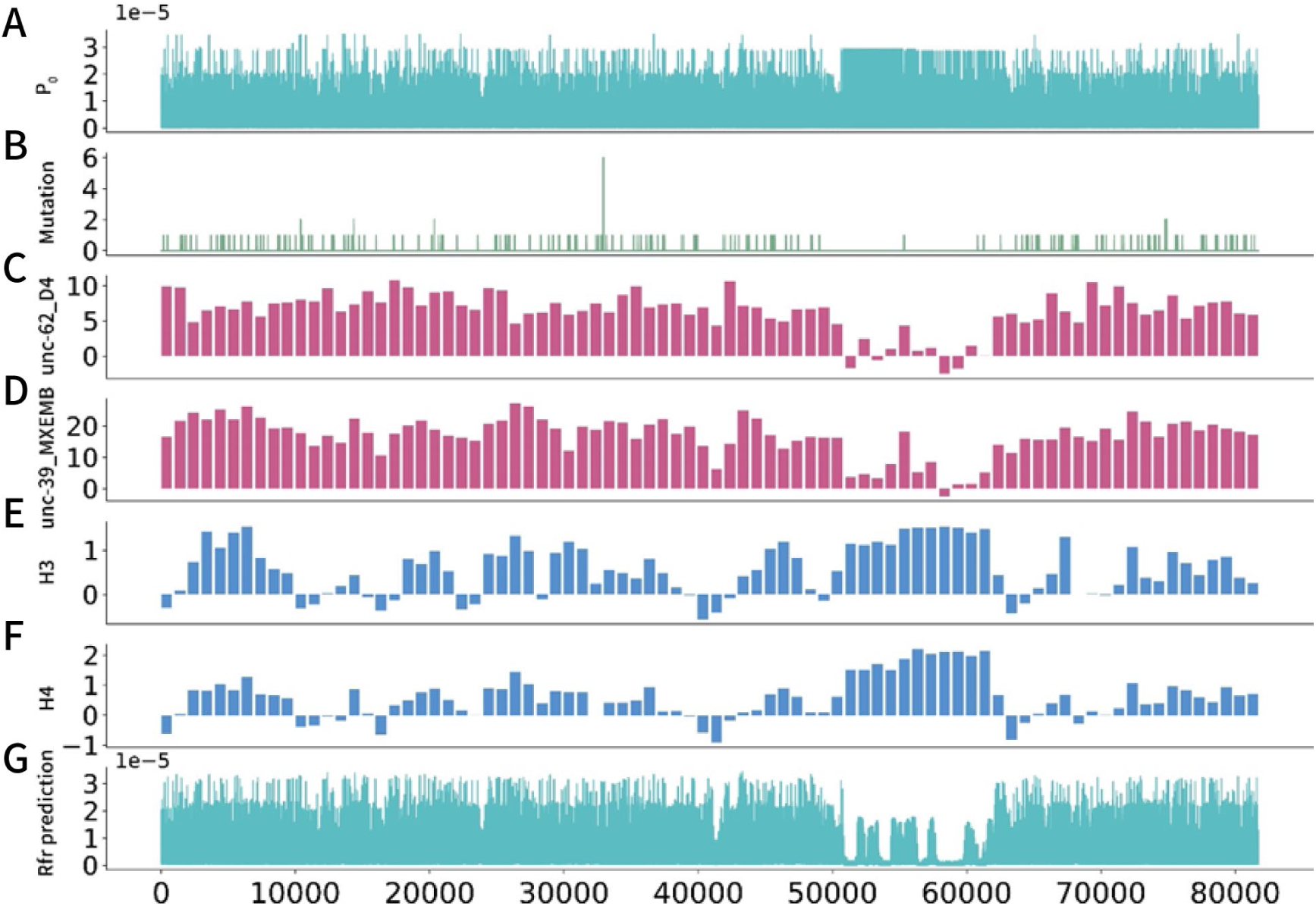
Mutation numbers in the MMP data shown association with the DNA-binding protein features. Take the MMP data and DNA-binding protein features on Chr V:6120909∼6202632 (*ttn-1*) as an example 1) Line plots shows the prediction made from the Flanking sequence preferences of EMS mutagenesis (*P*_0_). 2) The mutations observed in the MMP dataset. 3) Raw CHIP-chip signal data of unc-62 binding in young adult worms. 4) Raw CHIP-chip signal data of unc-39 binding in worm embryo. 5) Raw CHIP-seq signal data of histone H3 in L3 larval worms. 6) Raw CHIP-seq signal data of histone H4 in L3 larval worms. 7) Line plots shows the prediction made by a Random Forest regressor trained by DNA-binding protein data.

**Figure S3.**
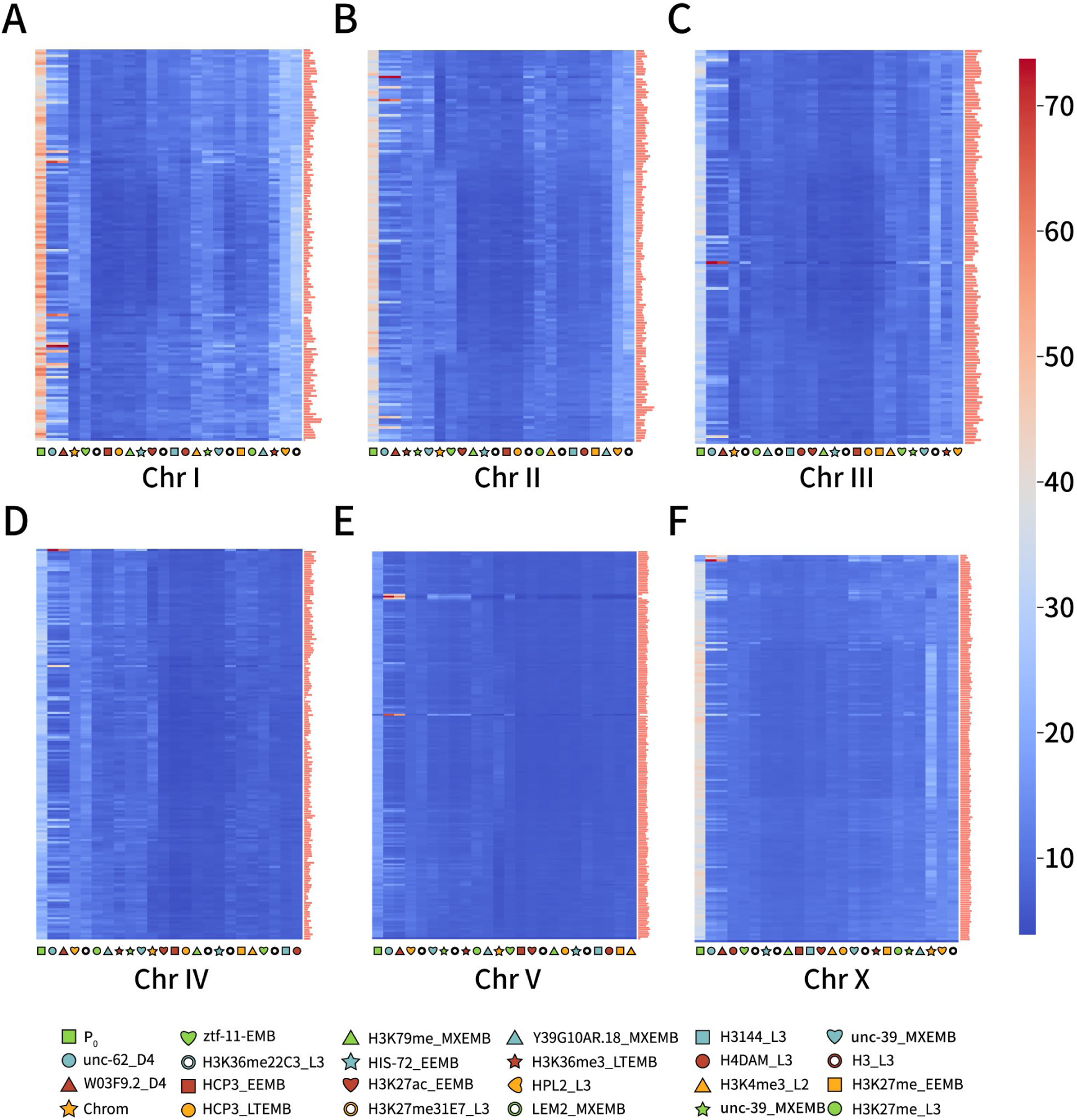
Feature importance analysis of the Random Forest regressor. A-F) Permutation importance (see Materials and Methods) of each features used to model the Random Forest regressor. The importance is shown in the heatmap alongside the chromosome. Mutation rate is normalized to 0∼1 and is shown in the bar plot attaching to the right side of the heatmap.

**Table S1.**

| **Feature Name** | **Information** | **Resource** |
| --- | --- | --- |
| Chrom | Which chromosome the base belongs to | Reference genome WS235 |
| Code | $P_0$ or 'pentabase bias' | Calculated with the MMP dataset |
| HPL2_L3 | ChIP-chip of L3 worm targeting heterochromatin protein HPL2; strain: N2 | GSE40947 |
| LEM2_MXEMB | ChIP-chip of mixed-stage embryo targeting nuclear envelope protein LEM2; strain: N2 | GSM562786 |
| unc-62_D4 | ChIP-seq of day4 young adult worm targeting transcription factor unc-62; strain: OP600 | GSM942047 |
| W03F9.2_D4 | ChIP-seq of day4 young adult worm targeting transcription factor W03F9.2; strain: OP215 | GSM729307 |
| H3K36me22C3_L3 | ChIP-chip of early embryo targeting K36trimethylated histone H3; strain: N2 | GSE22719 |
| HCP3_EEMB | ChIP-chip of early embryo targeting holocentric chromosome binding protein HCP3; strain: N2 | GSM1255286 |
| H3K27ac_EEMB | ChIP-chip of early embryo targeting K27 acetylated histone H3; strain: N2 | GSM562779 |
| HCP3_LTEMB | ChIP-chip of late embryo targeting holocentric chromosome binding protein HCP3; strain: N2 | GSM1255289 |
| H3K79me_MXEMB | ChIP-chip of mixed-stage embryo K79 trimethylated histone H3; strain: N2 | GSM562779 |
| HIS-72_EEMB | DNA tiling array of mixed-stage embryo; strain: JJ2061 | www.modencode.org 5326 |
| H3_L3 | ChIP-chip of L3 worm targeting histone H3; strain: N2 | GSM1005484 |
| H3K27me31E7_L3 | ChIP-chip of L3 worm targeting K27trimethylated histone H3; strain: N2 | GSM562735 |
| H3144_L3 | ChIP-chip of L3 worm targeting histone H3; strain: N2 | GSM562745 |
| H4DAM_L3 | ChIP-chip of L3 worm targeting histone H4; strain: N2 | GSM624427 |
| H3K36me3_LTEMB | ChIP-chip of late embryo targeting K36trimethylated histone H3; strain: N2 | GSM811293 |
| unc-62_EEMB | ChIP-seq of mixed-stage embryo targeting transcription factor unc-62; strain: OP600 | GSE48731 |
| unc-39_EEMB | ChIP-seq of mixed-stage embryo targeting transcription factor unc-39; strain: OP186 | GSE46775 |
| Y39G10AR.18_MXE MB | ChIP-chip of mixed-stage embryo targeting transcription factor Y39G10AR.18; strain: N2 | GSM811254 |
| H3K27me_L3 | ChIP-chip of L3 worm targeting K27 trimethylated histone H3; strain: N2 | GSM562735 |
| H3K27me_EEMB | ChIP-chip of early embryo targeting K27 trimethylated histone H3; strain: N2 | GSM811255 |
| H3K4me3_L2 | ChIP-chip of L2 worm targeting K4 trimethylated histone H3; strain: N2 | GSM811289 |
| ztf-11-EMB | ChIP-seq of mixed-stage embryo targeting zinc finger putative transcription factor ztf-11; strain: OP236 | GSE48746 |

**Table S2.** *C. elegans* Strains in this study.

*C. elegans* Strains in this study
| Strain Name | Genotype | Method |
| --- | --- | --- |
| N2 | *wild-type* | N.A |
| cas22599 | *OSM-3<sup>E251K</sup>* | Microinjection |
| cas23441 | [*OSM-3<sup>E251K</sup>*; *K04F10.2<sup>Arg506\*</sup>*] | Microinjection |

**Table S3.**
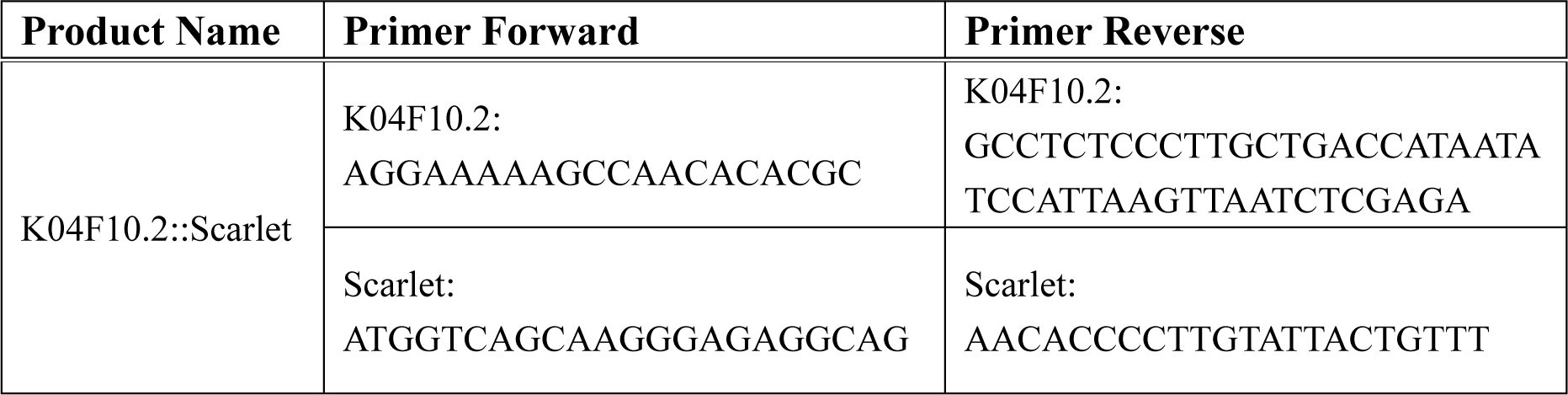
PCR products for *C. elegans* transgenesis.

## Notes

### Competing Interest Statement

The authors have declared no competing interest.

